# Nucleophagy contributes to genome stability though TOP2cc and nucleolar components degradation

**DOI:** 10.1101/2021.12.02.471011

**Authors:** Gabriel Muciño-Hernández, Adán Oswaldo Guerrero Cárdenas, Horacio Merchant-Larios, Susana Castro-Obregón

## Abstract

The nuclear architecture of mammalian cells can be altered as a consequence of anomalous accumulation of nuclear proteins or genomic alterations. Most of the knowledge about nuclear dynamics comes from studies on cancerous cells. How normal, healthy cells maintain genome stability avoiding accumulation of nuclear damaged material is less understood. Here we describe that primary mouse embryonic fibroblasts develop a basal level of nuclear buds and micronuclei, which increase after Etoposide-induced DNA Double-Stranded Breaks. These nuclear buds and micronuclei co-localize with autophagic proteins BECN1 and LC3 and with acidic vesicles, suggesting their clearance by nucleophagy. Some of the nuclear alterations also contain autophagic proteins and Type II DNA Topoisomerases (TOP2A and TOP2B), or nucleolar protein Fibrillarin, implying they are also targets of nucleophagy. We propose that a basal nucleophagy contributes to genome and nuclear stability and also in response to DNA damage and nucleolar stress.

## Introduction

Genome stability is essential for the proper function of the cells, and genome instability is a common feature of several pathologies primary affecting the nervous, immune and reproductive systems; genome instability also contributes to neurodevelopmental disorders, neurodegeneration, cancer development and premature aging (Ciccia and Elledge, 2010). Since early in development DNA is under constant endogenous challenges, for example when local abundant cell proliferation leads to DNA replication stress. Frequent genome dynamics has been uncovered recently, which occurs when cells produce DNA breaks as a physiological mechanism. For example, in response to TGFB1-induced epithelial-mesenchymal transition (Dobersch et al., 2021), as well as in active neurons (Madabhushi et al., 2015), DNA double-strand breaks (DSB) facilitate chromatin opening to initiate transcription of early-response genes. Another source of genome dynamics happens during the active recombination involving DSB that occurs during B and T immune cells differentiation to produce multiple antibodies and receptors, respectively. Interestingly, during neurogenesis perhaps a similar active recombination mediated by recurrent DSB also occurs, particularly in neuronal genes (Alt et al., 2017). This active DSB and repair could provide a mechanistic explanation to the mosaic nature of the mammalian brain recently described, pointing out to genome dynamics also in post-mitotic cells (Rohrback et al., 2018). Additionally, there are exogenous challenges to DNA integrity, such as DNA chemical modifications and DNA breaks caused by reactive oxygen species derived from normal cell metabolism, as well as exogenous sources, such as radiation and chemicals, giving rise to multiple types of DNA modifications. Therefore, DNA integrity needs to be constantly monitored and repaired throughout life. Eukaryotic cells have developed a network of intracellular pathways that sense DNA damage, signal to coordinate a cellular response and repair damaged DNA, collectively known as DNA damage response (DDR).

DDR regulates the recruitment of DNA repair molecules suitable to repair particular types of DNA damage. A defective DDR leads to genomic instability manifested either as minor chromosomal alterations or as pronounced chromosomal rearrangements (Langie et al., 2015). Due to the repetitive nature of rDNA present in the nucleolus, and its active transcription, this genomic region is especially susceptible to instability. Hence, chromosomal rearrangements of rDNA is frequently observed in tumors (Korsholm et al., 2020). Excessive accumulation of genomic alterations can affect the nuclear structure, eliciting the extrusion of nuclear content into the cytoplasm forming nuclear buds and micronuclei. The latter are fragments of chromosomes or whole chromosomes surrounded by nuclear envelop (Fenech et al., 2011; Kisurina-Evgenieva et al., 2016; Rao et al., 2008). The presence of micronuclei contributes to malignant cell transformation (Hintzsche et al., 2017). In senescent cells, fragments of chromatin containing damaged DNA are expelled from nuclei into the cytoplasm free of nuclear envelope, named cytoplasmic chromatin fragments (CFF) (Ivanov et al., 2013). Since micronuclei and nuclei-derived cytoplasmic material threaten genome integrity, it is essential to remove them.

Gene transcription, replication, recombination, etc. generate DNA entanglements (coiling and winding of the DNA double helix), which are resolved by DNA topoisomerases. Among them, Type II Topoisomerase (TOP2) catalyzes the resolution of DNA entanglements by creating transient DNA double-strand breaks that allow topological changes. During this process, TOP2 binds covalently to the 5’ end in the broken DNA forming a transitory intermediate cleavage complex (TOP2cc). Etoposide is a topoisomerase poison that stabilizes TOP2cc by misaligning DNA ends, preventing re-ligation, which results in trapping TOP2 on DNA termini, generating cytotoxic protein-linked DNA breaks that cells need to eliminate to avoid genome instability (Ashour et al., 2015; Austin et al., 2018).

In mammals, specifically in cancer cell lines undergoing genotoxic stress, macroautophagy (autophagy hereafter) is activated by genotoxic stress (Chen et al., 2015) and contributes to the removal of extruded nuclear material (Erenpreisa et al., 2011; Rello-Varona et al., 2012). Cytoplasmic chromatin fragments in senescent cells are also removed by autophagy (Ivanov et al., 2013). In autophagy deficient-cells chromosomal abnormalities and deficiencies in DNA damage repair occur (Bae and Guan, 2011; Chicote et al., 2020). Actually, autophagy seems to be protective of the genome, as the activation of different DNA repair pathways triggers autophagy, helping to resolve genomic instability (Eliopoulos et al., 2016).

The degradation of nuclear components by the autophagic machinery is coined Nucleophagy. Most of the data cited above has been obtained studying cancerous cells. In this work, we hypothesized that nucleophagy could be a mechanism to keep nuclear and genome integrity in normal (noncancerous) cells, in response to DNA damaging agents. We found that primary mouse embryonic fibroblasts (MEFs) developed nuclear buds and micronuclei in response to DSB caused by etoposide that contained damaged DNA and TOP2cc (Austin et al., 2018), as well as nucleolar components such as rRNA 2’-O-methyltransferase Fibrillarin. These nuclear alterations were surrounded by the autophagic proteins LC3 and BECN1, in proximity with lysosomal markers, indicative of their potential elimination by nucleophagy. Inhibition of autophagy reduced the frequency of nuclear alterations, suggesting an active role of the autophagic machinery in their formation. Interestingly, we observed a basal development of nuclear buds and micronuclei in healthy cells, which were also surrounded by nucleophagy machinery. Collectively, our data show that nucleophagy contributes to preserve nuclear cell physiology by constantly clearing damaged DNA through nuclear buds and micronuclei elimination, both basal and in response to genotoxic stress.

## Results

### There is a basal formation of nuclear buds and micronuclei in primary fibroblasts that increase with Etoposide-induced DSB

Since most of our knowledge about micronuclei formation and elimination comes from studies with cancerous cells, we aimed to study in primary cells micronuclei formation induced by DSB, which are the most toxic DNA lesions for cells. Primary MEFs were treated with 120 μM Etoposide for 2 h to cause DSB detectable by Neutral comet assay. To analyze DNA repair, Etoposide was removed. DSBs were gradually repaired becoming undetectable after 5h of recovery (Figure 1A-B and supplementary 1A). We also evaluated DDR activation by analyzing the recruitment of the phosphorylated histone γH2AX to sites with DSB. We observed at 2 h of Etoposide exposure abundant γH2AX, which was reduced after 5 h of recovery (figure 1C and S1B). Cell viability remained ≥80% during both DNA damage and repair (figure S1C). Large DNA damage leads to deformations of nuclear architecture and micronuclei formation (Kisurina-Evgenieva et al., 2016; Medvedeva et al., 2007), and the induction of multiple DSBs results in the budding of nuclear envelope and ultimately in micronuclei formation (Okamoto et al., 2012; Utani et al., 2011), hence, we analyzed the nuclear structure of MEFs treated with the sub-lethal dose of Etoposide. As expected, we found that Etoposide-treated cells bear nuclear protrusions or buds containing damaged DNA, identified by γH2AX (Figure 1D). We also observed cytoplasmic damaged DNA contained within micronuclei, as they were surrounded by lamin A/C and lamin B1. In some micronuclei we observed only lamin A/C (Figure 1E). Interestingly, we also found nuclear buds and micronuclei in a subpopulation of healthy, untreated cells, but lacking γH2AX, which suggest that these nuclear alterations contain a different kind of DNA damage (figure 1D). This observation implies that cells normally have a basal dynamic formation of nuclear buds and micronuclei, which to our knowledge has not been reported before. Interestingly, while the frequency of nuclear buds gradually increased after DNA damage and during DNA repair, the frequency of micronuclei also increased after DNA damage, but diminished upon DNA repair (figure 1D). These observations suggest that the dynamic of buds and micronuclei formation is similar, but micronuclei are being actively removed during DNA repair.

**Figure 1.**
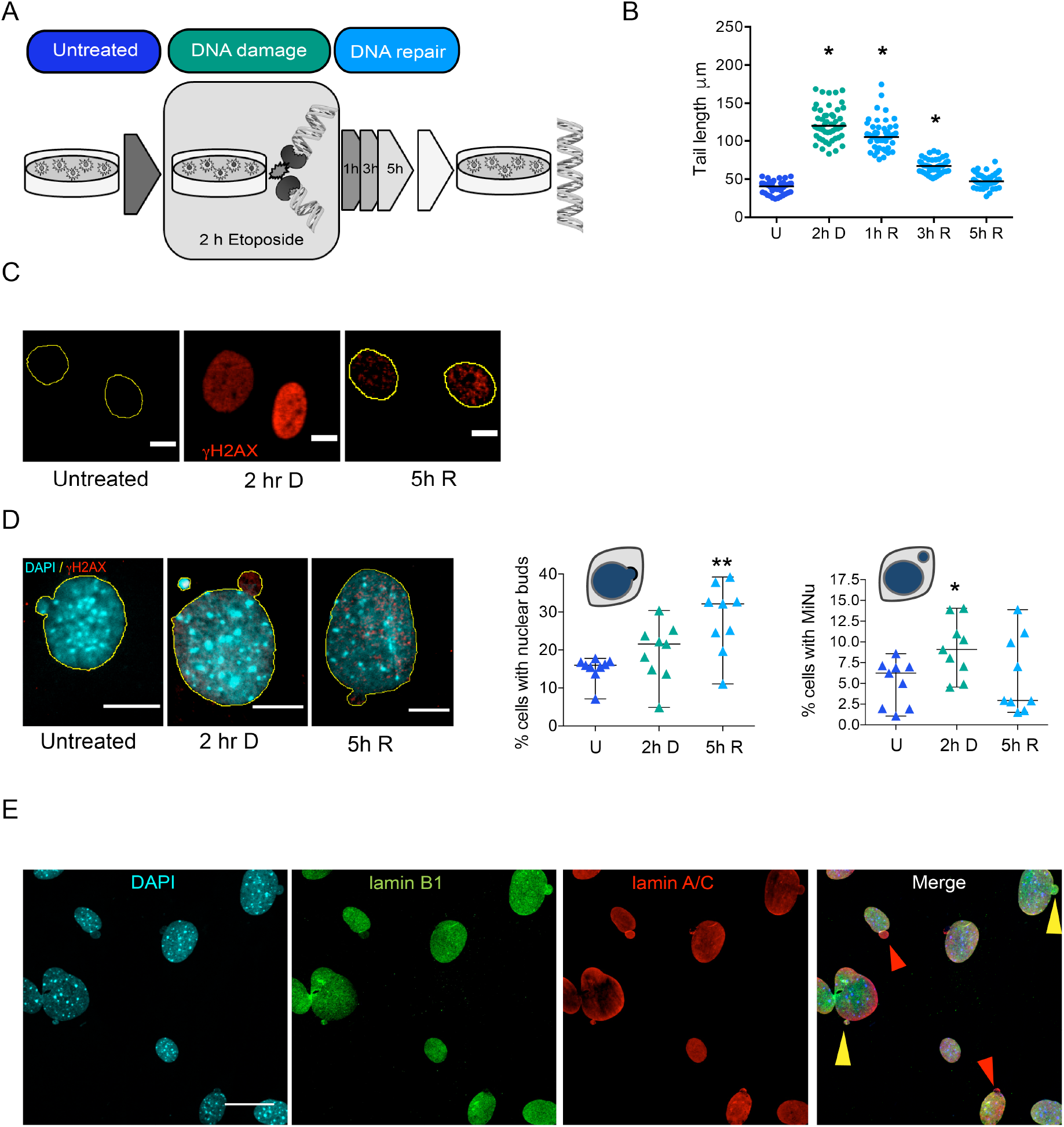
There is a basal formation of nuclear buds and micronuclei in primary fibroblasts, which increases with Etoposide-induced DBS. **A.** Workflow for DNA damage and repair assay. MEFs were exposed to Etoposide for 2 hours at 120 μM to damage DNA (2h D), then Etoposide was removed to allow DNA repair, which was monitored after 1, 3, and 5 h. **B.** Quantification of comet tails length (which is proportional to DSBs) in untreated cells (U), after 2 h of Etoposide exposure (2h D), and after 1h, 3h or 5hr of Etoposide removal (1h R, 3h R, 5h R, respectively). Bars represent median at each time point, with significant differences determined by Kruskal-Wallis test followed by Dunn’s multiple comparison test; *p<0.01. 50 comets were measured in each of three independent experiments. **C.** DDR followed by the recruitment of phospho-H2AX (γH2AX) in untreated (U), damaged (2h D) or repaired (5h R) DNA. Yellow contours indicate the nuclei of cells. Scale bars are equivalent to 10 μm. **D.** Nuclear buds or independent micronuclei (outlined in yellow) were observed by confocal microscopy in untreated (U), damaged (2h D) or repaired (5h R) DNA. DNA damaged marked with γH2AX (red) was found in both buds and micronuclei only when DNA was damaged. DNA was stained with DAPI. Scale bars are equivalent to 10 μm. Right, quantification of the percentage of cells with nuclear buds or micronuclei in untreated (U), damaged (2h D) or repaired (5h R) DNA. The mean ± SD of nine independent experiments is graphed, with significant differences determined by Mann Whitney test. For every experiment (represented as triangles) at least 50 cells were counted; * p<0.05, ** p<0.01. **E.** Representative immunofluorescence to detect Lamin-A/C (red) and Lamin-B1 (green) in MEFs treated with Etoposide for 2h (2h D). Yellow arrowheads show examples of buds containing both Lamina A/C and Lamina B1. Red arrowheads show examples of buds containing only Lamin A/C. Scale bar is equivalent to 30 μm.

### Nuclear buds and micronuclei are eliminated by nucleophagy in primary fibroblasts

In cancerous cell lines micronuclei removal is carried out by nucleophagy (Erenpreisa et al., 2011; Rello-Varona et al., 2012). We asked whether either basal or DNA damage-induced nuclear buds and micronuclei could also be eliminated by nucleophagy in primary MEFs.

We first analyzed the induction of the autophagy machinery in response to Etoposide. We monitored by Western blot the level of LC3-II, which results from LC3-I lipidation with phosphatidylethanolamine at the autophagosome membrane and its necessary for autophagosomal formation (Klionsky et al., 2021). Interestingly, we found two waves of autophagy activation, one upon DNA damage induction, increasing both LC3-I and LC3-II levels, which gradually decreased; a second slower induction of LC3-II formation occurred after one hour of DNA repair (1h after Etoposide removal), which was sustained along the repair process, and was reduced until DNA was completely repaired (5h) (Figures 2A and S2A). We then evaluated the potential role of the autophagic machinery in the elimination of nuclear buds and micronuclei. First, we followed the distribution of GFP-LC3 and found that more than 50% of the nuclear buds and micronuclei contained GFP-LC3 (figure 2B-C). We also monitored the intracellular distribution of BECN1, another protein required for autophagosome formation. Just as with LC3, we found nuclear buds and micronuclei containing BECN1 even in untreated cells, which tend to increase after DNA damage (figure 2B and 2D). We also noticed a nuclear enrichment of both GFP-LC3 and BECN1 (a representative wider field is shown in figure S2B), which agrees with previous observations indicating that autophagy mediates degradation of nuclear lamina through a direct interaction between LC3 and lamin B1 in proliferating cells. This interaction helps to translocate lamin B1 into the cytoplasm for its lysosomal degradation during oncogene-induced senescence (Dou et al., 2015). We speculated that LC3 could also contribute to the translocation of nuclear damaged material into the cytoplasm for autolysosomal degradation in primary cells. First, we confirmed the presence of LC3 in micronuclei, identified by containing of lamin-A/C (figure 2E). Then, we analyzed whether nuclear buds and micronuclei were associated with autolysosomes. We found micronuclei containing DNA, LC3 and stained with Lysotracker® (figure 2F, and with BECN1 in figure S2B). Interestingly, we noticed that a small percentage of untreated cells also had nuclear buds and micronuclei with autophagic components. This observation suggests that there is a basal level of nuclear dynamics, constantly forming nuclear protrusions and micronuclei, perhaps to eliminate genomic alterations that are frequently produced. To determine a causal role of nucleophagy for nuclear bus and micronuclei removal, the expression of *Atg7,* a member of the ubiquitin-like system required for autophagosome elongation (Simon et al., 2017), was silenced by iRNA previous to DNA damage induction. Surprisingly, the percentage of cells with nuclear buds decreased, both basal and caused by Etoposide-induced DSB (figure 2G). We also observed a subtle reduction in the percentage of cells with micronuclei (figure 2H). These results suggest that components of the autophagy machinery actively induce the formation of nuclear buds. In summary, we found nuclear buds and micronuclei with markers of different stages of the autophagic pathway, suggesting an active role of autophagy proteins in their formation, and a nucleophagic clearance of nuclear components.

**Figure 2.**
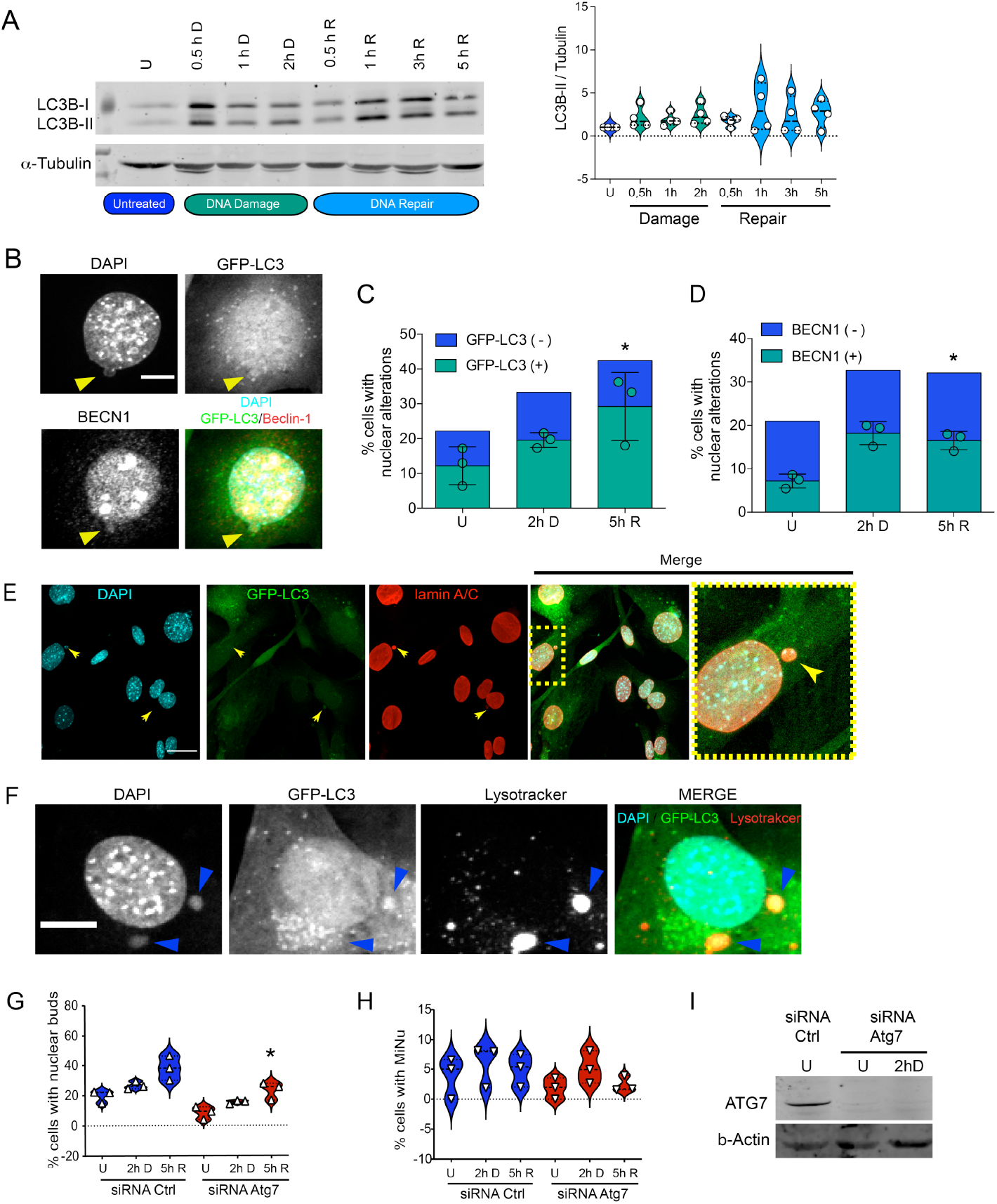
Nuclear buds and micronuclei are associated with components of different stages of the autophagic pathway. **A.** WB to detect LC3-I and LC3-II in MEFs untreated (U) or treated for 0.5, 1 and 2 h with Etoposide (DNA damage) and at 0.5, 1, 3 or 5 h after Etoposide removal (DNA repair). α-Tubulin was used as loading control. The graph on the right presents de distribution of LC3-II / α-Tubulin ratio obtained by densitometry of proteins detected by WB in four independent experiments. Even though there is a trend of LC3 induction upon Etoposide treatment, non-significant differences were obtained by ANOVA analysis followed by Kruskall-Wallis multiple comparison. **B.** Representative images of autophagic proteins GFP-LC3 and BECN1 found in nuclear buds (yellow arrowhead) in MEFs treated for 2 h with Etoposide used for quantifications shown in C and D. Scale bar represents to 10 μm. **C.** Percentage of cells with nuclear alterations (nuclear buds and micronuclei) is plotted. Of those, nuclear alterations containing GFP-LC3 are shown in green, without GFP-LC3 are shown in blue, from cells untreated (U), damaged (2h D) or with repaired DNA (5h R). **D.** Percentage of cells with nuclear alterations (nuclear buds and micronuclei) is plotted. Of those, nuclear alterations containing BECN1 are shown in green, without BECN1 and shown in blue, from cells untreated (U), damaged (2h D) or with repaired DNA (5h R). Color bars represent the mean of three independent experiments. Green symbols represent the percentage of cells with nuclear alterations containing GFP-LC3 or BECN1; bars represent ± SD. At least 50 cells were counted per treatment and experiment and significant differences were determined by Mann Whitney test *p<0.05. **E.** Representative micronuclei surrounded by lamin A/C containing GFP-LC3 (yellow arrows) in MEFs treated for 2 h with Etoposide. Yellow doted square indicates the magnified area shown to the right. Scale bar represents 30 μm. **F.** Representative micronuclei contained in autolysosomes, identified by having DNA, GFP-LC3 and Lysotracker® staining (blue arrowheads) in MEFs treated for 2 h with Etoposide. Scale bar corresponds to 10 μm. **G-I.** Functional autophagy seems to be necessary to form nuclear buds and micronuclei. MEF’s were transfected with siRNA control or siRNA *Atg7* for 48 h and then treated or not with Etoposide for 2 h and let repair DNA for 5 h (untreated (U), damaged (2h D) or repaired (5hR) DNA). Graphs show the percentage of cells with nuclear buds (**G**) or micronuclei (**H**). For every experiment at least 50 cells were counted by detecting DAPI signal in nuclear alterations in confocal images. The distribution of the data from three independent experiments is graphed. Significant differences were obtained after Two-way ANOVA analysis, followed by a Tukeýs multiple comparison test * p< 0.5 (5h R siRNA *Atg7 vs.* 5hR siRNA control). **I.** Western blot to verify *Atg7* silencing; β-actin was used as loading control. Whole blots are shown in supplemented figure S2C.

### Nucleophagy clears topoisomerase cleavage complex and nucleolar Fibrillarin

Several mechanisms to remove TOP2cc have been observed. For example, TOP2 can be disjoint by the protein TDP2 after partial TOP2 degradation. In an alternative mechanism, nucleases remove TOP2 and a fragment of DNA (Ashour et al., 2015; Pommier et al., 2016). We propose that nucleophagy may also contribute to the elimination of these complexes. To analyze it, we studied whether TOP2 were found outside the nucleus and within autophagosomes. By immunolocalization, we found both TOP2A and TOP2B in extra-nuclear bodies containing DNA at a basal level in healthy cells, which increased by Etoposide-induced DSB. We quantified the percentage of cells with cytoplasmic DNA, and around half of them contained TOP2A or TOP2B (Figure 3A-E). We found TOP2A within GFP-LC3 positive micronuclei (Figure 3A), and TOP2B within micronuclei also containing BECN1 (figure 3B). We confirmed by super-resolution microscopy TOP2B localization within micronuclei containing both DNA and BECN1 (figure 3C) in most of the micronuclei (figure 3F). After Etoposide-induced DSB we observed an increase in the frequency of micronuclei containing BECN1 and TOP2B, which then were reduced after DNA repair (figures S3B, S3F and S3H). We further demonstrated TOP2B nucleophagic degradation by immunogold co-localization of LC3 and TOP2B observed by transmission electron microscopy. TOP2B was found surrounded by LC3 in transit towards the cytoplasm, confirming the frequent nucleophagic degradation of nuclear alterations (Figure 3G).

**Figure 3.**
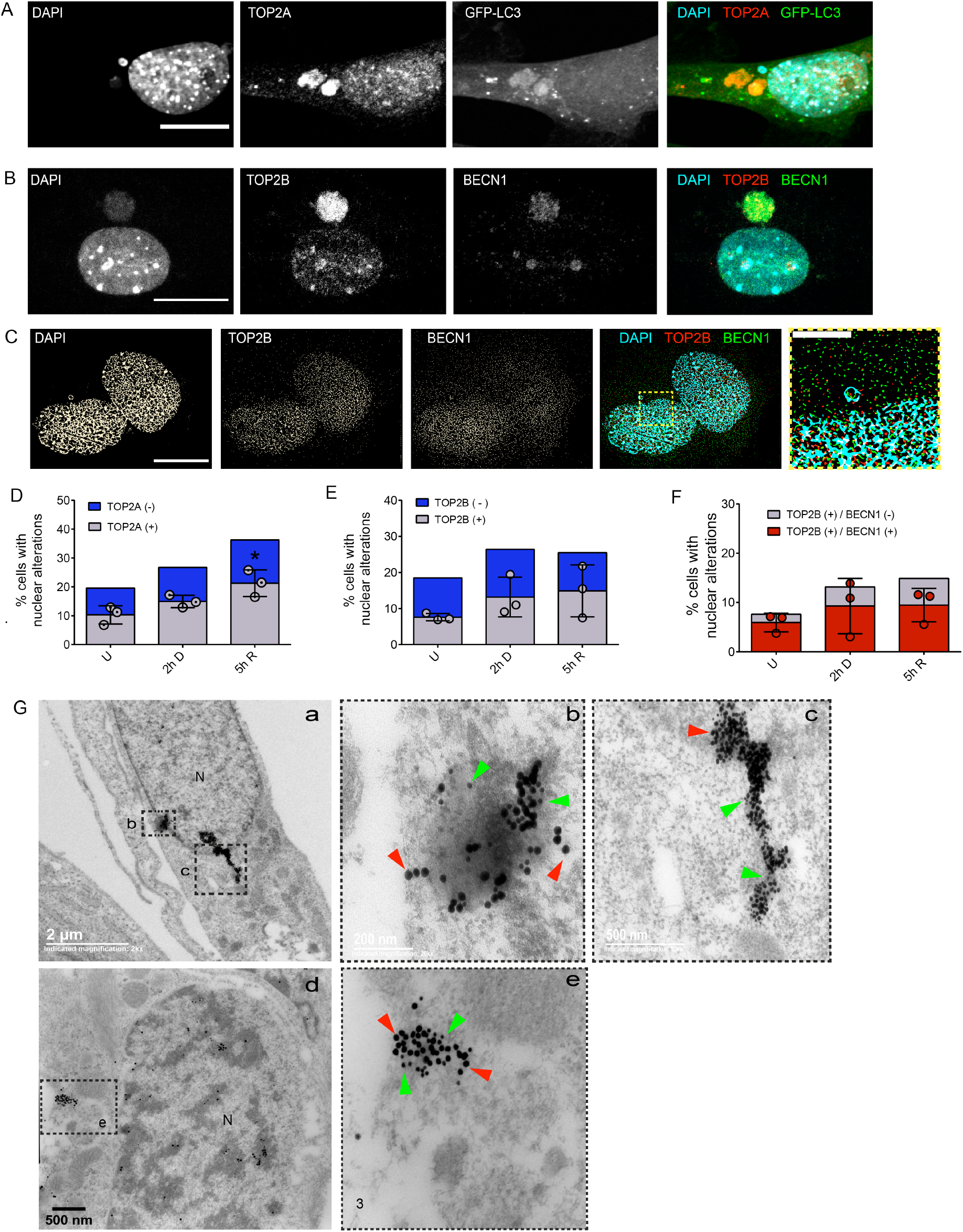
TOP2cc are targeted for nucleophagic clearance. **A.** Representative confocal image of an immunofluorescence to detect TOP2A in MEFs expressing GFP-LC3, treated with 120 μM Etoposide for 2h. Scale bar represents 20 μm. **B.** Representative confocal image of an immunofluorescence to detect TOP2B and BECN1 in MEFs, treated with 120 μM Etoposide for 2h. Scale bar represents 20 μm. **C.** Representative images obtained by super-resolution microscopy to detect co-localization of DNA and TOP2B (TOP2Bcc) with BECN1 in MEFs after 5h of DNA repair (yellow arrows). Yellow square represents the amplified section presented to the right. Scale bar represents 15 μm. **D.** Percentage of untreated (U), DNA damaged (2h D) or DNA repaired (5h R) cells with nuclear alterations (nuclear buds and micronuclei) containing DNA and TOP2A (grey bars). Nuclear alterations without TOP2A are shown in blue bars. The mean ± SD for three independent experiments (counting at least 50 cells per experiment) are graphed. (*) p<0.05, Mann Whitney test only for nuclear alterations with TOP2A. **E - F** Percentage of cells with nuclear alterations (nuclear buds and micronuclei) containing TOP2B (in **E**) or TOP2B co-localizing with Beclin1 (in **F**) in untreated MEFs or after DNA damage (2D, cells treated with 120 μM Etoposide for 2h) or DNA repair phase (5R, cells after 5h of Etoposide removal). At least 50 cells were counted for each experiment. The mean ± SD of three independent experiments is graphed. Non-significant differences were obtained after Kruskal-Wallis test. **G.** Electron micrographs showing simultaneous detection of LC3 and TOP2B by immunogold. Figures b and c indicate higher magnification areas of **a**, while **d** correspond to the lower magnification shown on figure **e**. Green arrows show examples of 15 nm gold particles coupled to secondary antibody to detect TOP2B and red arrows point to 25 nm gold particles coupled to secondary antibody to detect LC3.

To maintain genome stability in the ribosomal DNA domain is particularly challenging, since it is located in the nucleolus, a subnuclear compartment that has a high density of nucleic acids and proteins that creates a distinct environment that limits the accessibility of DNA repair factors (Korsholm et al., 2020). We considered that nucleosomal damage could also be removed by expelling nucleolar damaged material into the cytoplasm to be a nucleophagy target. In teratocarcinoma cells nucleolar aggresomes increase in response to chronic Etoposide exposure, and are transported to the cytoplasm where they are surrounded by the autophagic machinery (Salmina et al., 2017). We looked for the presence of Fibrillarin, a nucleolar marker, in micronuclei and nuclear buds in primary MEFs, treated or not with Etoposide. As shown in figure 4, Fibrillarin was found in micronuclei and nuclear buds in roughly 5% of the untreated cells, indicating a low basal level of nucleolar material exclusion from the nucleus. We confirmed by super-resolution microscopy the micronuclear nature of the cytoplasmic Fibrillarin, finding it with DNA surrounded by Lamin-A/C (figure 4B). As in previous experiments, close to 30% of the cells had nuclear alterations (nuclear buds and micronuclei); of those, around 16% of total nuclear alterations contained nucleolar material. In cells treated with etoposide we observed a mild increase of fibrillarin-containing nuclear buds and micronuclei, which remained in the same proportion of cells after 5 h of DNA repair. We then analyzed whether the nuclear buds and micronuclei containing Fibrillarin were also removed by nucelophagy. We detected GFP-LC3 in more than 50% of the nuclear alterations containing Fibrillarin in both untreated and etoposide-induced DBS cells. Noticeable, after 5 h of DNA repair most of the Fibrillarin-containing nuclear perturbation had GFP-LC3 (figures 4C). The presence of TOP2 and Fibrillarin in buds containing LC3 suggest that LC3 contributes to nucleolar bodies exiting the nucleus, driving them to nucleophagic degradation.

**Figure 4.**
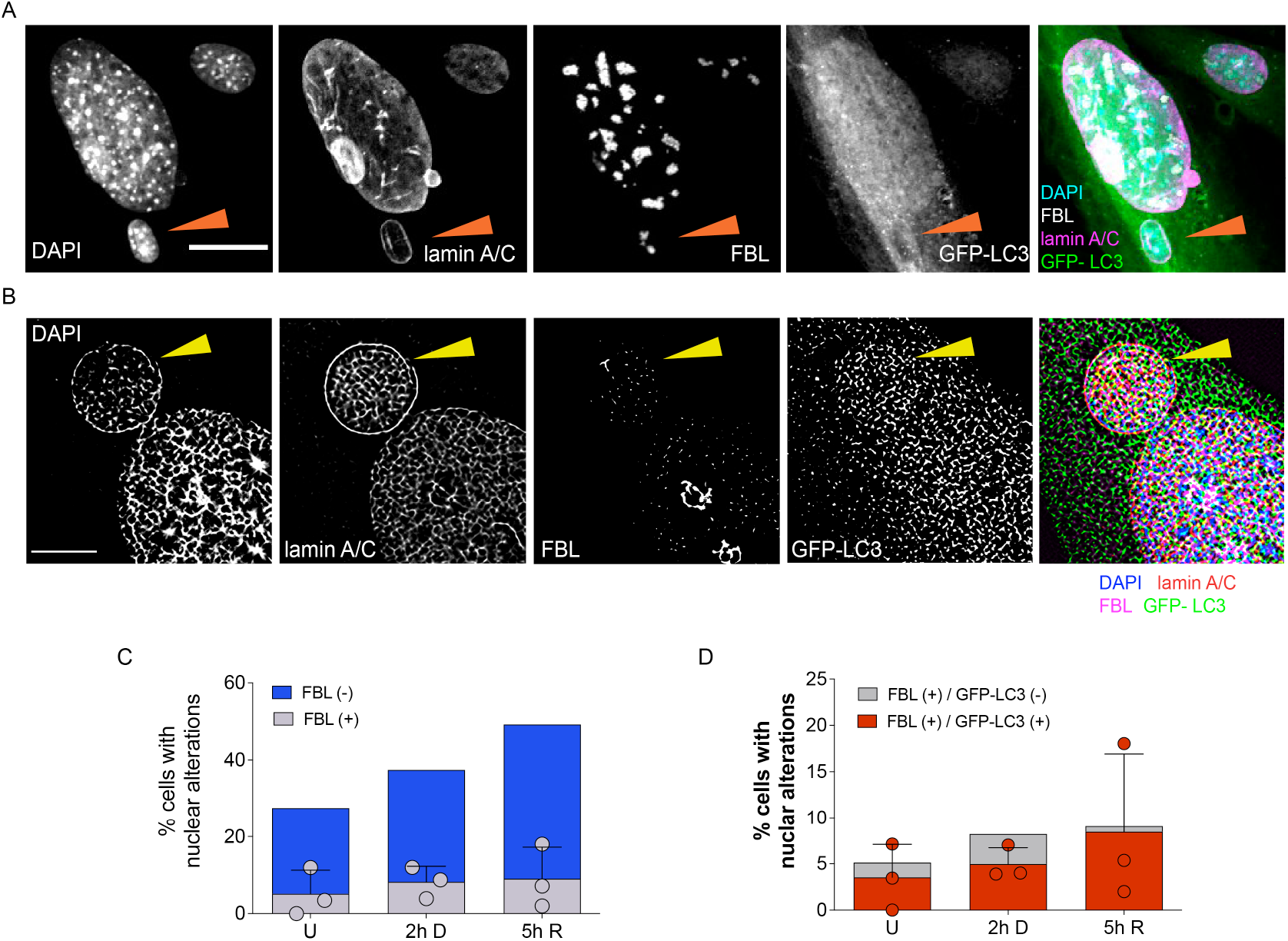
The nucleolus is a target for nucleophagic degradation. **A,** Representative confocal microscopy image showing the nucleolar protein Fibrillarin in a micronucleus containing DNA stained with DAPI and surrounded by lamin-A/C (arrow head). This micronucleus also has the autophagic marker GFP-LC3, in a cell treated with etoposide for 2 hr. Scale bar represents 20 μm. **B,** Representative super-resolution microscopy images of the same experiment described in **A**. Scale bar represents 5 μm. **C.** Percentage of cells containing nuclear alterations (buds and micronuclei), with (grey bars) or without (blue bars) Fibrillarin in untreated (U), Etoposide-treated (2h D) or after 5 hr of DNA repair (5h R). **D.** Percentage of cells containing nuclear alterations (buds and micronuclei), with (grey bars) or without (red bars) Fibrillarin and GFP-LC3 in untreated (U), Etoposide-treated (2h D) or after 5 hr of DNA repair (5h R). Dots represent the mean of each experiment (n=3); at least 50 cells were counted per experiment by analyzing DAPI distribution in confocal images. Bars correspond to SD. Even though there is a trend to increase nuclear alterations, non-significant differences were obtained after Kruskal-Wallis test followed by Dunn’s multiple comparison test.

## Discussion

Several studies have shown that autophagy contributes to genome stability by different mechanisms, for example it elevates the level of DNA repair proteins of both HR and NHEJ pathways, and enhances DNA damage recognition to be repaired by nucleotide excision repair (Lin et al., 2015; Liu et al., 2015). Additionally, BECN1 interacts directly with TOP2B, which leads to the activation of DNA repair proteins, and the formation of NR and DNA-PK repair complexes (Xu et al., 2017). Cytoplasmic elimination of nuclear lamina components has been observed in cells with oncogenic insults, with LC3 interacting directly with both lamin-associated domains on chromatin and Lamin-B1 and Lamin A/C to be targeted for lysosomal degradation (Dou et al., 2015; Lenain et al., 2015; Li et al., 2019). These observations lead to the conclusion that nucleophagy contributes to tumor suppression. Here we describe that BECN1, together with LC3, could have a pivotal role in cells with Etoposide-induced DSB integrating the DNA repair machinery with the autophagy machinery, promoting the nuclear extrusion of damaged DNA, TOP2cc and Fibrillarin through nuclear buds and micronuclei formation, that would later be eliminated by nucleophagy. Interestingly, we observed also a basal formation and elimination of nuclear buds and micronuclei. Both Etoposide-induced and basal buds and micronuclei formation depend on ATG7, as silencing its expression reduced their abundance (Figure 2G-H). This observation supports the contribution of autophagy machinery to extrude nuclear damaged material.

We noticed that the basal formation of micronuclei did not have γH2AX, suggesting that the nuclear material to be extruded does not contain DSB but other type of damaged DNA, or perhaps that they are formed by a proteostasis dysfunction not involving DNA damage. The proteasome also degrades TOP2 (Mao et al., 2001) and Fibrillarin (Chen et al., 2002). Their co-localization with autophagic proteins in nuclear alterations suggests nucleophagy has a collaborative with the proteasome, protecting both genome integrity and nuclear morphology. The recruitment of multiple molecules for DNA repair into the nucleus could trigger an imbalance in proteostasis by overloading the proteasome. The resulting accumulation of molecules could affect nuclear structure, which is necessary to avoid.

An outcome of the overloaded activity of UPS is the accumulation and aggregation of polyubiquitinated proteins as aggresomes (Latonen et al., 2011). This occurs in the cytoplasm but also in the nucleoplasm, specifically at nucleoli, where under different stress conditions (heat shock, acidosis or genotoxic insults) proteins, RNA and conjugated Ubiquitin accumulate (Jacob et al., 2013; Latonen, 2011; Latonen, 2019) For example, under DNA damage, it occurs an early and transient nucleolar accumulation of paraspeckle proteins (Moore et al., 2011) (paraspeckles are nuclear subcompartments which function as a reservoir for splicing factors (Nunes and Moretti, 2017)), and E2F1, affecting the structure and function of the nucleolus (Jin et al., 2014). The final destiny for aggresomes is not totally understood, but it has been suggested their persistence until the recovery of UPS degradative capacity to clear their components (Latonen, 2011). Another possibility is that the aggresome is cleared by autophagy to promote genome and cell viability (Salmina et al., 2017). Accordingly, we observed some nuclear alterations in MEFs containing Fibrillarin, which tend to increase during DNA damage and repair. A proportion of such nuclear alterations, mainly nuclear buds, bear nuclear lamina proteins and GFP-LC3, suggesting the formation of autophagic vesicles with nucleolar components to be targeted to lysosomes. Hence, nucleophagy could be a mechanism to alleviate also nucleolar stress.

Along the results presented here, the damage inflicted by Etoposide detected by a wide γH2AX signal, implies a huge quantity of TOP2cc that has to be removed. We propose that nucleophagy supports the proteasome, phosphodiesterases and nucleases in the removal of peptides or whole TOP2 protein, free and complexed with DNA, that accumulates into the nucleus during DNA damage and repair. Despite topoisomerases have not been reported as targets for autophagic degradation, Alchanati *et al*., reported a decrease in TOP2A levels when cancerous cells were grown under glucose deprivation (Alchanati et al., 2009), an autophagy-inducing condition (Klionsky et al., 2016).

By a careful analysis of the protrusions and micronuclei formed, we observed a variable composition on their envelope. In some cases we detected only one type of nuclear lamina, either Lamin-A/C or Lamin-B1, while in other cases, both nuclear laminas were present (figure 2B). The different micronuclear envelope composition is probably related to different DNA damage or different DNA structures affected that lead to their formation. Others have also identified structural differences in micronuclei envelope with variable presence of Lamin B1, which has been linked to different abilities to replicate (Okamoto et al., 2012) or repair (Terradas et al., 2012) the genome, by affecting the recruitment of proteins required in those processes. These differences in the envelope composition could affect their nucleophagic clrearance.

In summary, as depicted in figure 5, we propose that Etoposide-induced DSBs in MEFs, in addition to lead to an accumulation of TOP2cc, also promotes aggresomes formation in the nucleolus, and both events contribute to nuclear distortions, causing genotoxic stress. To contend with such stress, proteasome and nucelophagy function in a dynamic way inside the nucleus. Insufficiencies on any of these homeostatic systems imply a risk of genomic instability, which in turn could drive the cell into a senescent or malignant state.

**Figure 5.**
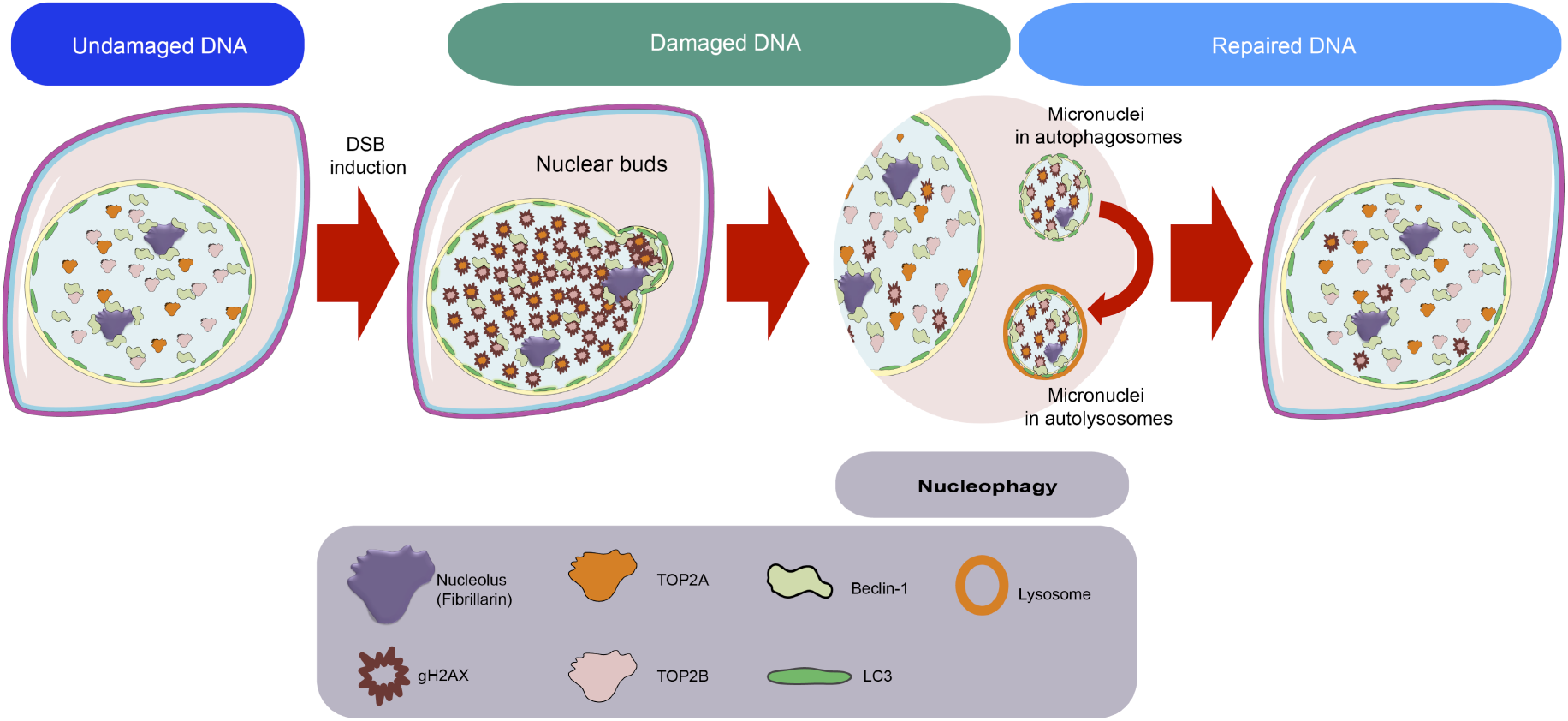
Integrative model. Intensive DNA damage alter the cell physiology, especially inside the nucleus, the capacity of systems to eliminate macromolecules are surpassed promoting the accumulation of nucleic acids and proteins which leads to nuclear deformations and ultimately to the expulsion of nuclear material to the cytoplasm. This process is facilitated by autophagic machinery, which at the end culminates with the elimination of the nuclear damaged material in the lysosome.

It is important to take into account that autophagy has different roles in cancer development and is affected differently by chemotherapeutic drugs. Interfering with autophagic function might alter nuclear dynamics and maintenance.

## Material and methods

### Cells and culture conditions

All the experiments were done with mouse embryonic fibroblasts (MEFs) at cell passage 4 or 5. MEFs from wild type CD1 or GFP-LC3 transgenic mice (C57BL/6J) (Mizushima et al., 2004) were obtained at E13.5 (for wild type mice) or E13 (for transgenic GFP-LC3 mice) according to the standard protocol (Xu, 2005). Animals were obtained from the animal house of the Institute of Cellular Physiology (IFC) at the National Autonomous University of Mexico (UNAM), housed at 22 ^0^C in 12h light/12h dark cycle with *ad libitum* access to water and food. Mice used in the present study were handled and cared according to the animal care and ethics legislation. All procedures were approved by the Internal Committee of Care and Use of Laboratory Animals of the IFC (IFC-SCO174-21). No contamination of mycoplasma was tested in every batch using the VenorGeM Mycoplasma Detection Kit (SIGMA-Aldrich MP-0025, St.Louis MO, USA), following the procedure indicated by the provider. MEFs were grown in Dulbeccós Modified Eagle Medium + GlutaMAXTM, 10% FBS and 100 U/mL Penicillin/Streptomycin. Media and supplements were from GIBCO® Life TechnologiesTM, Grand Island, NY, USA. Culture conditions consisted of a humidified and 5% CO2 atmosphere at 37° C. DNA damage was induced by incubating cells with Etoposide (Etopos® injectable solution, Lemery, Mexico City) at 120 uM for 0.5, 1 and 2 hours. Then Etoposide was removed and cells were washed twice with PBS 1X and incubated for 1, 3, 5 or 24 h in fresh medium.

#### siRNA Transfection

WT MEFs were transfected using Lipofectamine 2000 (Invitrogen, Carlsbad, CA, USA) according to manufacturer instructions. Briefly, 5×10^4^ cells/well were seeded into 12 well plates 24 hr before transfection and using antibiotic-free medium. For each well, 20 pmol siRNA and 3 µl Lipofectamine were mixed and added for 6 h. After that fresh antibiotic-free medium was added and cells were incubated for 48 h. SMARTpool siRNA ATG7-FITC from Dharmacon (Lafayette, CO, USA). Control siRNA targets a region of a Luciferase coding gene.

### Neutral comet assay

The DSB were detected using a neutral comet assay. Briefly about 100 cells/µL were resuspended in PBS and mixed at a 1:5 ratio with 0.% low-melting point agarose (BIO RAD Certified™ Low Melt Agarose #1613112, Hercules, California, USA) at 37°C. Then with the help of a coverslip, about 50 μL of the previous mix were spread on glass slides pre-coated with 1% normal-melting point agarose (BIO RAD Certified™ PCR Agarose #1613104, Hercules, California, USA). The slides were incubated first at 4° C for 2 min and then for an extra 10 min at room temperature. After the removal of coverslip each slide was sequentially covered and incubated for 60 min with pre-chilled lysis solution and then with unwinding buffer at 4°C. Next electrophoresis ran with slides by applying 25 V for 20 min. After that slides were incubated in neutralization buffer for 10 min, repeating this steps 3 times. At the end with SYBR green (solution 1:10000 in PBS 1X, SYBRTM green I Nucleic Acid Gel Stain, INVITROGEN, Eugene, Oregon, USA) DNA was stained. Lysis solution: 0.03 M EDTA, 1% SDS. Unwinding and electrophoresis buffer: Tris 60 mM, Acetic acid 90 mM, EDTA 2.5 mM, pH 9.0. Neutralization buffer: Tris-HCl 500 mM, pH 7.5

To visualize the comets (DNA), a NIKON ECLIPSE Ti-U microscope with 20X magnification and the NIS Elements BR software (Nikon Instruments Inc ®, NY, USA) images were acquired and analyzed. For analysis, length and area of broken DNA were determined by processing 50 comet images for each treatment.

### Immunofluorescence

The day before treatments, cells were grown on coverslips at a density of 2.5 x 10^4^ cells/cm^2^ on 12-well plates. After treatments, at room temperature cells were fixed with 4% paraformaldehyde for 30 min, then washed with PBS, permeabilized for 5 min with PBS / 0.5% Triton and blocked for 1h with PBS/4% BSA. Coverslips were incubated overnight at 4°C with primary antibody (diluted in PBS/2%BSA). The next day, later than removing the primary antibody and a wash with PBS, AlexaFluor-conjugated secondary antibodies (diluted 1:500 in PBS/2%BSA) (LIFE TECHNOLOGIES, Oregon, USA) were added and incubated for 1 h at room temperature. Finally nuclei were stained with DAPI (1 µg/mL) for 10 min.

Primary antibodies used for immunofluorescence: mouse anit-gamma-H2AX (1:1000, ABCAM ab26350, Cambridge, MA, USA), mouse anti-Lamin A/C (1:1000, SANTA CRUZ BIOTECHNOLOGY), rabbit anti-Lamin B1 (1:1000), rabbit anti-LC3B, rabbit anti-Beclin1 (1:100, SANTA CRUZ BIOTECHNOLOGY sc-11427), mouse anti-TOP2A (1:100, SANTA CRUZ BIOTECHNOLOGY sc-365916), mouse anti-TOP2B (1:100, SANTA CRUZ BIOTECHNOLOGY sc-25330), rabbit anti-Fibrillarin (1:1000, ABCAM ab5821, Cambridge, MA, USA).

Secondary antibodies: Goat anti-mouse IgG (H+L) Alexa Fluor A594 (1:500) (A11032), Goat anti-rabbit IgG (H+L) Alexa Fluor A594 (1:500) (A11037) Goat anti-mouse IgG (H+L) Alexa Fluor A488 (1:500) (A11029), Goat anti-rabbit IgG (H+L) Alexa Fluor A488 (1:500) (A11029) All secondary antibodies were from LIFE TECHNOLOGIES, Oregon, USA except Donkey anti-rabbit IgG (H+L) Alexa Fluor 647 which was from *Jackson Immuno Research Laboratories, Inc,* Pennsylvania, USA.

### Immunoblotting analysis

Using a buffer with 62.5 mM Tris (pH 6.5), 2% SDS and 2 mg/ml protease inhibitor 18 (Complete, ROCHE MOLECULAR DIAGNOSTICS, Pleasanton CA, USA) cells were lysed. Between 30 to 120 micrograms of protein lysates were separated by SDS-PAGE and then transferred to polyvinylidene fluoride (PVDF) membranes. Following a 1h blocking step, membranes were incubated overnight with primary antibodies. Secondary antibody IRDye® 680RD goat anti-rabbit (925-68071, LI-COR) or IRDye® 800CW goat anti-mouse (925-32210, LI-COR) were added at 1:5000 dilution in TTBS. Scan was done using the Odyssey® IR scanner, the image acquisition and analysis were on Odyssey® Image Studio software 5.2.5. Blocking solution consisted in 3% nonfat dry milk (Blotting-Grade Blocker, BIO-RAD Laboratories, Inc. USA, Cat. # 170-6404) in TTBS.

Primary antibodies: mouse anit-gamma-H2AX (1:1000 ABCAM 26350, Cambridge, MA, USA), rabbit anti-LC3B (1:1000), rabbit anti-ATG7 (1:1000), rabbit anti-βActin (1:10,000), mouse anti-αTubulin (1:10000, Cell Signaling 3873, Beverly, MA, USA).

### Confocal Imaging

All images were collected as Z-stacks with an LSM800 (Zeiss) confocal microscope using 40x/1.3 Oil immersion objective with 1 Airy unit aperture of pinhole. Samples were excited with 405 nm, 488 nm, 561nm and 640 nm laser lines. CZI files obtained with ZEISS ZEN software and images of Z-projection were processed in Fiji (imageJ) software.

### Immunoelectron microscopy

Cells were fixed with 3% glutaraldehyde. Following fixation, dehydration was performed in an ethanol gradient: 30-40-50-60-70-80-90-100 % ethanol at 4°C. Then, the cells were embedded in a LR White resin and polymerization was carried out at 50 °C. Ultrathin sections of 70-80 nm were cut from the polymer using an Ultracut-Recheirt-Jung and placed on nickel grids for immunogold assay.

The thin sections were washed twice for 2 min with deionized water and two times with PBS with 0.005 % Tween20. Sections were then incubated for 30 min with the blocking solution (50 mM glycine, 0.005 % Tween20, 0.01 % Triton X-100 and 0.1 % BSA in PBS)(Rosas-Arellano et al., 2016). After blocking, sections were incubated with the primary antibody: rabbit anti-LC3 (1:500, MBL PD014, Nagoya, Japan). After rinsing three times in PBS with 0.005 % Tween20, the sections were incubated overnight at 4 °C with the secondary antibody (1:20). Samples were washed three times with PBS, 0.005 % Tween20 and post-fixed in 2 % glutaraldehyde in PBS for 10 min. The sections were then rinsed with distilled water twice for 5 min and contrasted with 2 % uranyl acetate, rinsed with water, dried and observed under a JEOL JEM 1200 EXII electron microscope.

Secondary antibodies: donkey anti-rabbit IgG (H&L) conjugated with 25-nm gold particles (Electron Microscopy Science Aurion #25708, PA, USA), donkey anti-mouse IgG (H&L) conjugated with 15-nm gold particles (Electron Microscopy Science Aurion #25817, PA, USA).

### Super resolution microscopy

Super resolution microscopy imaging was performed at the National Laboratory for Advanced Microcopy (LNMA) of UNAM. Immunofluorescence samples were imaged on a Nanoimager-S (Oxford Nanoimaging Ltd) using widefield fluorescence excitation. Samples were excited by alternating laser illumination with a 405 (DAPI), 473 (Alexa Fluor A488, GFP) and 561 (Alexa Fluor A594) laser lines. Detection of the signal was achieved via an 100X, 1.4 NA, oil-immersion objective (Olympus) and an sCMOS Hamamatsu Orca Flash 4.0 V3 using an embedded image splitter for dual-channel fluorescence acquisition. Imaging time = 33 ms, effective pixel size at object plane = 117 nm. Sub diffraction images were obtained via SRRF, a multi-frame super-resolution microscopy approach which gathers nanoscopic information from the statistical analysis of sequences of images collected at the same imaging plane [https://www.nature.com/articles/ncomms12471]. Each super resolution image was derived by the analysis of serial stacks of 100 images collected per fluorescence excitation channel. Each serial stack was drift corrected and analyzed using the NanoJ-core and NanoJ-SRRF plugins of FIJI / Image J [https://iopscience.iop.org/article/10.1088/1361-6463/ab0261]. Parameters used for SRRF computation were ring radius 0.5, radiality magnification 10, axes in ring 8 parameters, Temporal Analysis: AVG. The rest of the parameters were left as the recommended default values [https://iopscience.iop.org/article/10.1088/1361-6463/ab0261].

### Statistical analysis

Graphs and data analysis were done with GraphPad Prism 6 (GraphPad Software Inc. La Jolla, CA, USA). Different test to determine statistically significant differences were applied as indicated in every figure

## Acknowledgements

We are thankful to Dr. Beatriz Aguilar for her technical assistance. We are thankful to Dr. Ruth Rincón for confocal analysis assistance, to M.Sc. Rodolfo Paredes for electron microscopy imaging and to Dr. Abraham Rosas for both confocal analysis and electron microscopy imaging, all at the Imagenology Unit at IFC; we also thank to José P. Oviedo for his assistance for super resolution imaging at LNMA. We are thankful to Claudia Rivero at the Animal Facility, to M.C. Ana Maria Escalante and Francisco Pérez at the IT Unit, a to Aurey Galvan and Manuel Ortínez at the Equipment Maintenance Workshop, all at IFC. Data in this work are part of the doctoral dissertation in the “Programa de Doctorado en Ciencias Bioquímicas” at the Universidad Nacional Autónoma de México (UNAM) of GMH.

## Authors contribution

GMH conceived and designed the work, performed the experiments, interpreted the results and drafted the first version of the manuscript. AOGC performed the super resolution microscopy analysis and interpreted the results; HML acquired electron microscopy images and interpreted the results; SCO conceived and designed the work, interpreted the results and revised the manuscript. All authors approved the final version of the manuscript and are accountable for all aspects of the work.

## Supplementary Figures

**Figure S1.**
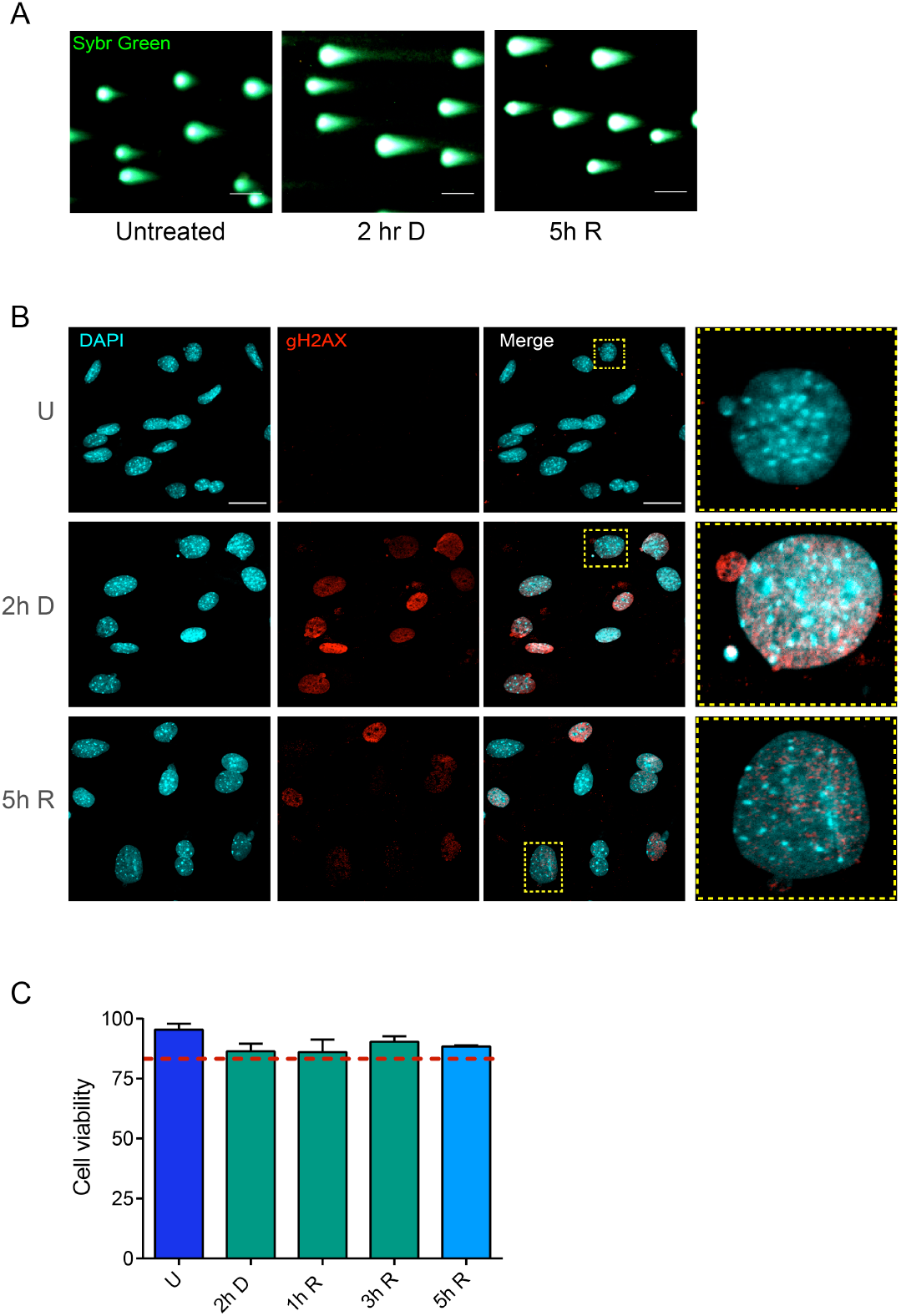
Etoposide treatment in primary MEFs causes DBS, DDR response and increases nuclear alterations. **A.** Representative images of Comet assay used to quantify data resented in Figure 1b, without treatment (Undamaged), after 2h of Etoposide treatment (2 h D) and after DNA repair (5 h R). DNA was stained with Sybr Green®. Scale bar is equivalent to 100 μm. **B.** DDR was monitored by the immunodetection of γH2AX (in red) in the nuclei of cells at the same time points as in A. At every stage (undamaged, damaged and repaired DNA) there are nuclear alterations observed as nuclear buds or cytoplasmic micronuclei containing DNA marked with γH2AX. DNA was stained with DAPI. Scale bar is equivalent to 30 μm. **C.** 2h of Etoposide treatment is sub-lethal. Cell viability was determined by Trypan blue exclusion in MEFs treated or not (U) with Etoposide for 2 hr (2h D) or at the indicated times after Etoposide removal (1hR, 3h R, 5h R). Data are presented as mean ± SD from three independent experiments. Red dashed line points 80% of the cell viability.

**Figure S2.**
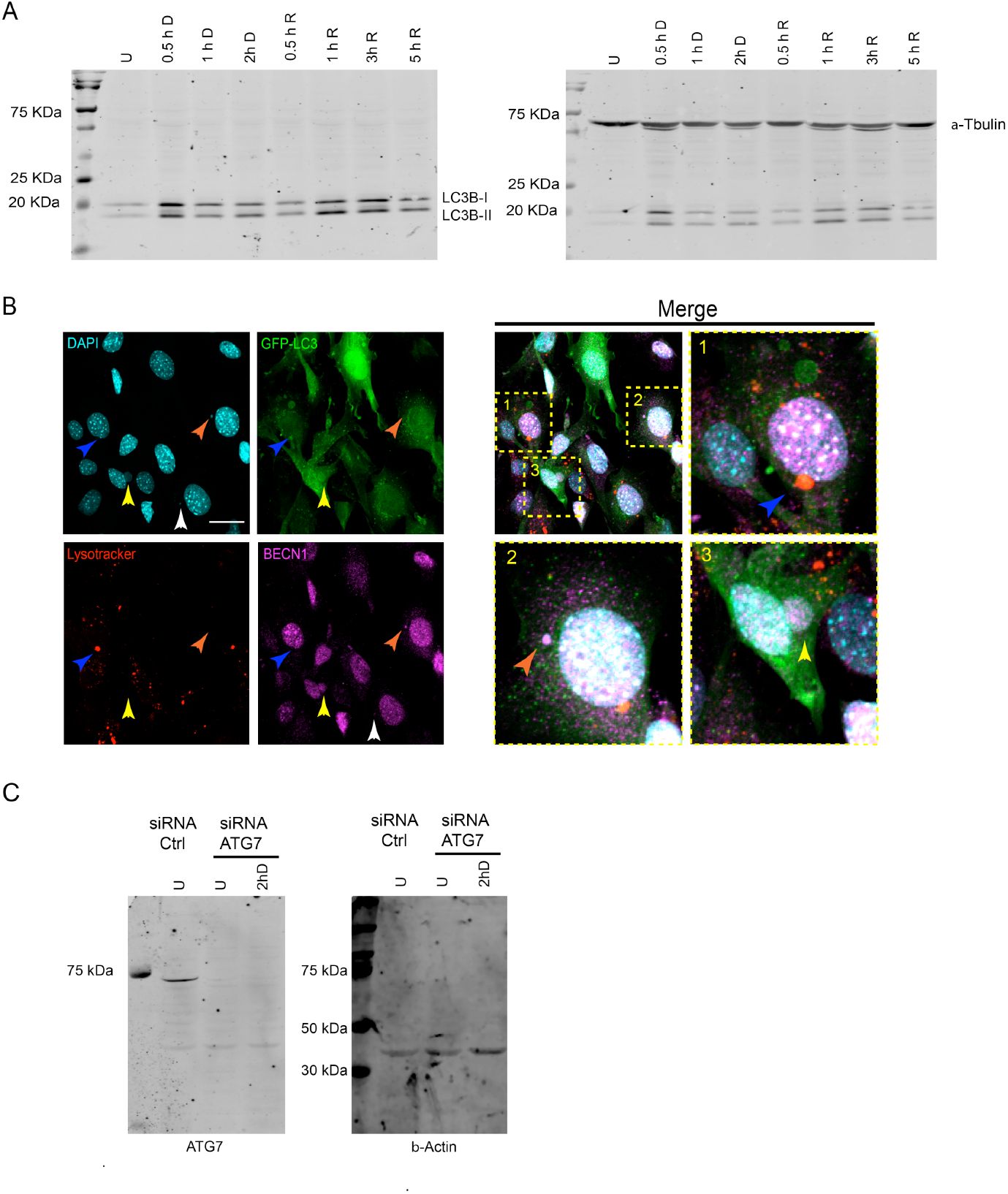
**A.** Whole blots of the WB shown in figure 2A to detect LC3-I, LC3-II and α-Tubulin. **B.** Some micronuclei look surrounded by autophagic markers and acidic vesicles. Orange arrows indicate co-localization between DNA and BECN1, yellow arrows show co-localization between DNA, GFP-LC3 and BECN1, and blue arrows indicate a micronucleus surrounded by GFP-LC3 and Lysotracker® (autolysosome). Yellow squares correspond to amplified sections. Cells were treated with Etoposide for 2h. Scale bars correspond to 30 μm. **C**. Whole membranes for WB to detect ATG7 shown in Figure 2I. MEFs were transfected with siRNA-Atg7 or control siRNA for 48h and then untreated or treated with Etoposide for 2h to detect before total protein extraction. β-actin was detected as loading control.

**Figure S3.**
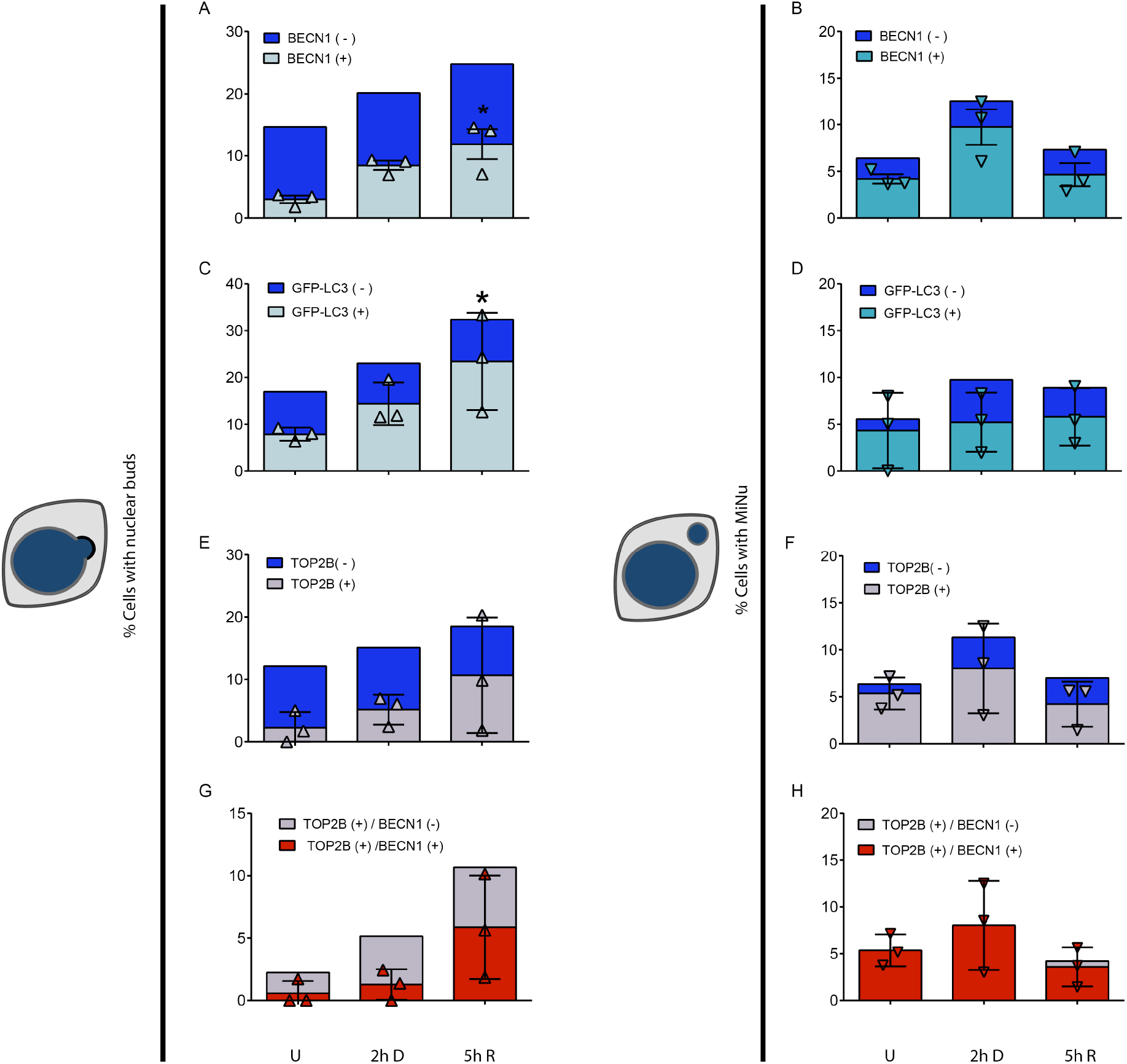
Nuclear alterations contain autophagic markers and TOP2B. **A, B, C and D.** Complimentary graphs showing the differential quantification of nuclear buds and micronuclei containing Beclin-1 (figures A and B) or LC3 (GFP-LC3 in figures C and D) or TOP2B (figures E and F). **G and H**. Complimentary quantifications of nuclear alterations containing simultaneously Beclin-1 and TOP2B. The mean of three independent experiments are graphed. Only for the indicated proteins are presented the result for every experiment as symbols (triangles and inverted triangles) with bars representing SD. Just for buds containing GFP-LC3, (*) p<0.05, Kruskal-Wallis test followed by Dunn’s multiple comparison test.

## References

Alchanati, I., C. Teicher, G. Cohen, V. Shemesh, H.M. Barr, P. Nakache, D. Ben-Avraham, A. Idelevich, I. Angel, N. Livnah, S. Tuvia, Y. Reiss, D. Taglicht, and O. Erez. 2009. The E3 ubiquitin-ligase Bmi1/Ring1A controls the proteasomal degradation of Top2alpha cleavage complex - a potentially new drug target. PLoS One. 4:e8104.

Alt, F.W., P.C. Wei, and B. Schwer. 2017. Recurrently Breaking Genes in Neural Progenitors: Potential Roles of DNA Breaks in Neuronal Function, Degeneration and Cancer. *In* Genome Editing in Neurosciences. R. Jaenisch, F. Zhang, and F. Gage, editors, Cham (CH). 63–72.

Ashour, M.E., R. Atteya, and S.F. El-Khamisy. 2015. Topoisomerase-mediated chromosomal break repair: an emerging player in many games. Nat Rev Cancer. 15:137–151.

Austin, C.A., K.C. Lee, R.L. Swan, M.M. Khazeem, C.M. Manville, P. Cridland, A. Treumann, A. Porter, N.J. Morris, and I.G. Cowell. 2018. TOP2B: The First Thirty Years. Int J Mol Sci. 19.

Bae, H., and J.L. Guan. 2011. Suppression of autophagy by FIP200 deletion impairs DNA damage repair and increases cell death upon treatments with anticancer agents. Mol Cancer Res. 9:1232–1241.

Chen, J.H., P. Zhang, W.D. Chen, D.D. Li, X.Q. Wu, R. Deng, L. Jiao, X. Li, J. Ji, G.K. Feng, Y.X. Zeng, J.W. Jiang, and X.F. Zhu. 2015. ATM-mediated PTEN phosphorylation promotes PTEN nuclear translocation and autophagy in response to DNA-damaging agents in cancer cells. Autophagy. 11:239–252.

Chen, M., T. Rockel, G. Steinweger, P. Hemmerich, J. Risch, and A. von Mikecz. 2002. Subcellular recruitment of fibrillarin to nucleoplasmic proteasomes: implications for processing of a nucleolar autoantigen. Mol Biol Cell. 13:3576–3587.

Chicote, J., V.J. Yuste, J. Boix, and J. Ribas. 2020. Cell Death Triggered by the Autophagy Inhibitory Drug 3-Methyladenine in Growing Conditions Proceeds With DNA Damage. Front Pharmacol. 11:580343.

Ciccia, A., and S.J. Elledge. 2010. The DNA damage response: making it safe to play with knives. Mol Cell. 40:179–204.

Dobersch, S., K. Rubio, I. Singh, S. Gunther, J. Graumann, J. Cordero, R. Castillo-Negrete, M.B. Huynh, A. Mehta, P. Braubach, H. Cabrera-Fuentes, J. Bernhagen, C.M. Chao, S. Bellusci, A. Gunther, K.T. Preissner, S. Kugel, G. Dobreva, M. Wygrecka, T. Braun, D. Papy-Garcia, and G. Barreto. 2021. Positioning of nucleosomes containing gamma-H2AX precedes active DNA demethylation and transcription initiation. Nat Commun. 12:1072.

Dou, Z., C. Xu, G. Donahue, T. Shimi, J.A. Pan, J. Zhu, A. Ivanov, B.C. Capell, A.M. Drake, P.P. Shah, J.M. Catanzaro, M.D. Ricketts, T. Lamark, S.A. Adam, R. Marmorstein, W.X. Zong, T. Johansen, R.D. Goldman, P.D. Adams, and S.L. Berger. 2015. Autophagy mediates degradation of nuclear lamina. Nature. 527:105–109.

Eliopoulos, A.G., S. Havaki, and V.G. Gorgoulis. 2016. DNA Damage Response and Autophagy: A Meaningful Partnership. Front Genet. 7:204.

Erenpreisa, J., K. Salmina, A. Huna, E.A. Kosmacek, M.S. Cragg, F. Ianzini, and A.P. Anisimov. 2011. Polyploid tumour cells elicit paradiploid progeny through depolyploidizing divisions and regulated autophagic degradation. Cell Biol Int. 35:687–695.

Fenech, M., M. Kirsch-Volders, A.T. Natarajan, J. Surralles, J.W. Crott, J. Parry, H. Norppa, D.A. Eastmond, J.D. Tucker, and P. Thomas. 2011. Molecular mechanisms of micronucleus, nucleoplasmic bridge and nuclear bud formation in mammalian and human cells. Mutagenesis. 26:125–132.

Hintzsche, H., U. Hemmann, A. Poth, D. Utesch, J. Lott, H. Stopper, and G.f.U.-M. Working Group “In vitro micronucleus test. 2017. Fate of micronuclei and micronucleated cells. Mutat Res. 771:85–98.

Ivanov, A., J. Pawlikowski, I. Manoharan, J. van Tuyn, D.M. Nelson, T.S. Rai, P.P. Shah, G. Hewitt, V.I. Korolchuk, J.F. Passos, H. Wu, S.L. Berger, and P.D. Adams. 2013. Lysosome-mediated processing of chromatin in senescence. J Cell Biol. 202:129–143.

Jacob, M.D., T.E. Audas, J. Uniacke, L. Trinkle-Mulcahy, and S. Lee. 2013. Environmental cues induce a long noncoding RNA-dependent remodeling of the nucleolus. Mol Biol Cell. 24:2943–2953.

Jin, Y.Q., G.S. An, J.H. Ni, S.Y. Li, and H.T. Jia. 2014. ATM-dependent E2F1 accumulation in the nucleolus is an indicator of ribosomal stress in early response to DNA damage. Cell Cycle. 13:1627–1638.

Kisurina-Evgenieva, O.P., O.I. Sutiagina, and G.E. Onishchenko. 2016. Biogenesis of Micronuclei. Biochemistry (Mosc). 81:453–464.

Klionsky, D.J., A.K. Abdel-Aziz, S. Abdelfatah, M. Abdellatif, A. Abdoli, S. Abel, H. Abeliovich, M.H. Abildgaard, Y.P. Abudu, A. Acevedo-Arozena, I.E. Adamopoulos, K. Adeli, T.E. Adolph, A. Adornetto, E. Aflaki, G. Agam, A. Agarwal, B.B. Aggarwal, M. Agnello, P. Agostinis, J.N. Agrewala, A. Agrotis, P.V. Aguilar, S.T. Ahmad, Z.M. Ahmed, U. Ahumada-Castro, S. Aits, S. Aizawa, Y. Akkoc, T. Akoumianaki, H.A. Akpinar, A.M. Al-Abd, L. Al-Akra, A. Al-Gharaibeh, M.A. Alaoui-Jamali, S. Alberti, E. Alcocer-Gomez, C. Alessandri, M. Ali, M.A. Alim Al-Bari, S. Aliwaini, J. Alizadeh, E. Almacellas, A. Almasan, A. Alonso, G.D. Alonso, N. Altan-Bonnet, D.C. Altieri, E.M.C. Alvarez, S. Alves, C. Alves da Costa, M.M. Alzaharna, M. Amadio, C. Amantini, C. Amaral, S. Ambrosio, A.O. Amer, V. Ammanathan, Z. An, S.U. Andersen, S.A. Andrabi, M. Andrade-Silva, A.M. Andres, S. Angelini, D. Ann, U.C. Anozie, M.Y. Ansari, P. Antas, A. Antebi, Z. Anton, T. Anwar, L. Apetoh, N. Apostolova, T. Araki, Y. Araki, K. Arasaki, W.L. Araujo, J. Araya, C. Arden, M.A. Arevalo, S. Arguelles, E. Arias, J. Arikkath, H. Arimoto, A.R. Ariosa, D. Armstrong-James, L. Arnaune-Pelloquin, A. Aroca, D.S. Arroyo, I. Arsov, R. Artero, D.M.L. Asaro, M. Aschner, M. Ashrafizadeh, O. Ashur-Fabian, A.G. Atanasov, A.K. Au, P. Auberger, H.W. Auner, L. Aurelian, et al. 2021. Guidelines for the use and interpretation of assays for monitoring autophagy (4th edition)(1). Autophagy. 17:1–382.

Klionsky, D.J., K. Abdelmohsen, A. Abe, M.J. Abedin, H. Abeliovich, A. Acevedo Arozena, H. Adachi, C.M. Adams, P.D. Adams, K. Adeli, P.J. Adhihetty, S.G. Adler, G. Agam, R. Agarwal, M.K. Aghi, M. Agnello, P. Agostinis, P.V. Aguilar, J. Aguirre-Ghiso, E.M. Airoldi, S. Ait-Si-Ali, T. Akematsu, E.T. Akporiaye, M. Al-Rubeai, G.M. Albaiceta, C. Albanese, D. Albani, M.L. Albert, J. Aldudo, H. Algul, M. Alirezaei, I. Alloza, A. Almasan, M. Almonte-Beceril, E.S. Alnemri, C. Alonso, N. Altan-Bonnet, D.C. Altieri, S. Alvarez, L. Alvarez-Erviti, S. Alves, G. Amadoro, A. Amano, C. Amantini, S. Ambrosio, I. Amelio, A.O. Amer, M. Amessou, A. Amon, Z. An, F.A. Anania, S.U. Andersen, U.P. Andley, C.K. Andreadi, N. Andrieu-Abadie, A. Anel, D.K. Ann, S. Anoopkumar-Dukie, M. Antonioli, H. Aoki, N. Apostolova, S. Aquila, K. Aquilano, K. Araki, E. Arama, A. Aranda, J. Araya, A. Arcaro, E. Arias, H. Arimoto, A.R. Ariosa, J.L. Armstrong, T. Arnould, I. Arsov, K. Asanuma, V. Askanas, E. Asselin, R. Atarashi, S.S. Atherton, J.D. Atkin, L.D. Attardi, P. Auberger, G. Auburger, L. Aurelian, R. Autelli, L. Avagliano, M.L. Avantaggiati, L. Avrahami, S. Awale, N. Azad, T. Bachetti, J.M. Backer, D.H. Bae, J.S. Bae, O.N. Bae, S.H. Bae, E.H. Baehrecke, S.H. Baek, S. Baghdiguian, A. Bagniewska-Zadworna, et al. 2016. Guidelines for the use and interpretation of assays for monitoring autophagy (3rd edition). Autophagy. 12:1–222.

Korsholm, L.M., Z. Gal, B. Nieto, O. Quevedo, S. Boukoura, C.C. Lund, and D.H. Larsen. 2020. Recent advances in the nucleolar responses to DNA double-strand breaks. Nucleic Acids Res. 48:9449–9461.

Langie, S.A., G. Koppen, D. Desaulniers, F. Al-Mulla, R. Al-Temaimi, A. Amedei, A. Azqueta, W.H. Bisson, D.G. Brown, G. Brunborg, A.K. Charles, T. Chen, A. Colacci, F. Darroudi, S. Forte, L. Gonzalez, R.A. Hamid, L.E. Knudsen, L. Leyns, A. Lopez de Cerain Salsamendi, L. Memeo, C. Mondello, C. Mothersill, A.K. Olsen, S. Pavanello, J. Raju, E. Rojas, R. Roy, E.P. Ryan, P. Ostrosky-Wegman, H.K. Salem, A.I. Scovassi, N. Singh, M. Vaccari, F.J. Van Schooten, M. Valverde, J. Woodrick, L. Zhang, N. van Larebeke, M. Kirsch-Volders, and A.R. Collins. 2015. Causes of genome instability: the effect of low dose chemical exposures in modern society. Carcinogenesis. 36 Suppl 1:S61–88.

Latonen, L. 2011. Nucleolar aggresomes as counterparts of cytoplasmic aggresomes in proteotoxic stress. Proteasome inhibitors induce nuclear ribonucleoprotein inclusions that accumulate several key factors of neurodegenerative diseases and cancer. Bioessays. 33:386–395.

Latonen, L. 2019. Phase-to-Phase With Nucleoli -Stress Responses, Protein Aggregation and Novel Roles of RNA. Front Cell Neurosci. 13:151.

Latonen, L., H.M. Moore, B. Bai, S. Jaamaa, and M. Laiho. 2011. Proteasome inhibitors induce nucleolar aggregation of proteasome target proteins and polyadenylated RNA by altering ubiquitin availability. Oncogene. 30:790–805.

Lin, W., N. Yuan, Z. Wang, Y. Cao, Y. Fang, X. Li, F. Xu, L. Song, J. Wang, H. Zhang, L. Yan, L. Xu, X. Zhang, S. Zhang, and J. Wang. 2015. Autophagy confers DNA damage repair pathways to protect the hematopoietic system from nuclear radiation injury. Sci Rep. 5:12362.

Liu, E.Y., N. Xu, J. O’Prey, L.Y. Lao, S. Joshi, J.S. Long, M. O’Prey, D.R. Croft, F. Beaumatin, A.D. Baudot, M. Mrschtik, M. Rosenfeldt, Y. Zhang, D.A. Gillespie, and K.M. Ryan. 2015. Loss of autophagy causes a synthetic lethal deficiency in DNA repair. Proc Natl Acad Sci U S A. 112:773–778.

Madabhushi, R., F. Gao, A.R. Pfenning, L. Pan, S. Yamakawa, J. Seo, R. Rueda, T.X. Phan, H. Yamakawa, P.C. Pao, R.T. Stott, E. Gjoneska, A. Nott, S. Cho, M. Kellis, and L.H. Tsai. 2015. Activity-Induced DNA Breaks Govern the Expression of Neuronal Early-Response Genes. Cell. 161:1592–1605.

Mao, Y., S.D. Desai, C.Y. Ting, J. Hwang, and L.F. Liu. 2001. 26 S proteasome-mediated degradation of topoisomerase II cleavable complexes. J Biol Chem. 276:40652–40658.

Medvedeva, N.G., I.V. Panyutin, I.G. Panyutin, and R.D. Neumann. 2007. Phosphorylation of histone H2AX in radiation-induced micronuclei. Radiat Res. 168:493–498.

Mizushima, N., A. Yamamoto, M. Matsui, T. Yoshimori, and Y. Ohsumi. 2004. In vivo analysis of autophagy in response to nutrient starvation using transgenic mice expressing a fluorescent autophagosome marker. Mol Biol Cell. 15:1101–1111.

Moore, H.M., B. Bai, F.M. Boisvert, L. Latonen, V. Rantanen, J.C. Simpson, R. Pepperkok, A.I. Lamond, and M. Laiho. 2011. Quantitative proteomics and dynamic imaging of the nucleolus reveal distinct responses to UV and ionizing radiation. Mol Cell Proteomics. 10:M111 009241.

Nunes, V.S., and N.S. Moretti. 2017. Nuclear subcompartments: an overview. Cell Biol Int. 41:2–7.

Okamoto, A., K. Utani, and N. Shimizu. 2012. DNA replication occurs in all lamina positive micronuclei, but never in lamina negative micronuclei. Mutagenesis. 27:323–327.

Pommier, Y., Y. Sun, S.N. Huang, and J.L. Nitiss. 2016. Roles of eukaryotic topoisomerases in transcription, replication and genomic stability. Nat Rev Mol Cell Biol. 17:703–721.

Rao, X., Y. Zhang, Q. Yi, H. Hou, B. Xu, L. Chu, Y. Huang, W. Zhang, M. Fenech, and Q. Shi. 2008. Multiple origins of spontaneously arising micronuclei in HeLa cells: direct evidence from long-term live cell imaging. Mutat Res. 646:41–49.

Rello-Varona, S., D. Lissa, S. Shen, M. Niso-Santano, L. Senovilla, G. Marino, I. Vitale, M. Jemaa, F. Harper, G. Pierron, M. Castedo, and G. Kroemer. 2012. Autophagic removal of micronuclei. Cell Cycle. 11:170–176.

Rohrback, S., B. Siddoway, C.S. Liu, and J. Chun. 2018. Genomic mosaicism in the developing and adult brain. Dev Neurobiol. 78:1026–1048.

Rosas-Arellano, A., J.B. Villalobos-Gonzalez, L. Palma-Tirado, F.A. Beltran, A. Carabez-Trejo, F. Missirlis, and M.A. Castro. 2016. A simple solution for antibody signal enhancement in immunofluorescence and triple immunogold assays. Histochem Cell Biol. 146:421–430.

Salmina, K., A. Huna, I. Inashkina, A. Belyayev, J. Krigerts, L. Pastova, A. Vazquez-Martin, and J. Erenpreisa. 2017. Nucleolar aggresomes mediate release of pericentric heterochromatin and nuclear destruction of genotoxically treated cancer cells. Nucleus. 8:205–221.

Simon, H.U., R. Friis, S.W. Tait, and K.M. Ryan. 2017. Retrograde signaling from autophagy modulates stress responses. Sci Signal. 10.

Terradas, M., M. Martin, L. Hernandez, L. Tusell, and A. Genesca. 2012. Nuclear envelope defects impede a proper response to micronuclear DNA lesions. Mutat Res. 729:35–40.

Utani, K., A. Okamoto, and N. Shimizu. 2011. Generation of micronuclei during interphase by coupling between cytoplasmic membrane blebbing and nuclear budding. PLoS One. 6:e27233.

Xu, F., Y. Fang, L. Yan, L. Xu, S. Zhang, Y. Cao, L. Xu, X. Zhang, J. Xie, G. Jiang, C. Ge, N. An, D. Zhou, N. Yuan, and J. Wang. 2017. Nuclear localization of Beclin 1 promotes radiation-induced DNA damage repair independent of autophagy. Sci Rep. 7:45385.

Xu, J. 2005. Preparation, culture, and immortalization of mouse embryonic fibroblasts. Curr Protoc Mol Biol. Chapter 28:Unit 28 21.

