## Supplementary material for "Nucleophagy contributes to genome stability though TOP2cc and nucleolar components degradation"

### Supplementary Figures

FIGURE S1

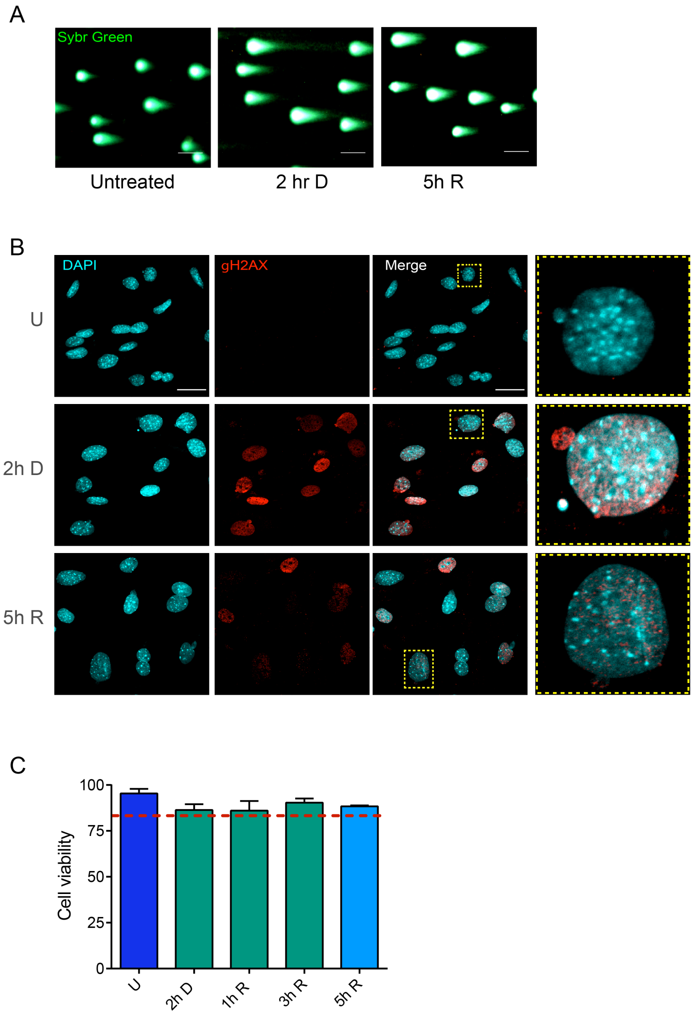


**Figure S1. Etoposide treatment in primary MEFs causes DBS, DDR response and increases nuclear alterations. A.** Representative images of Comet assay used to quantify data resented in Figure 1b, without treatment (Undamaged), after 2h of Etoposide treatment (2 h D) and after DNA repair (5 h R). DNA was stained with Sybr Green®. Scale bar is equivalent to 100 $\mu$m. **B.** DDR was monitored by the immunodetection of $\gamma$H2AX (in red) in the nuclei of cells at the same time points as in A. At every stage (undamaged, damaged and repaired DNA) there are nuclear alterations observed as nuclear buds or cytoplasmic micronuclei containing DNA marked with $\gamma$H2AX. DNA was stained with DAPI. Scale bar is equivalent to 30 $\mu$m. **C.** 2h of Etoposide treatment is sub-lethal. Cell viability was determined by Trypan blue exclusion in MEFs treated or not (U) with Etoposide for 2 hr (2h D) or at the indicated times after Etoposide removal (1hR, 3h R, 5h R). Data are presented as mean ± SD from three independent experiments. Red dashed line points 80% of the cell viability.

FIGURE S2

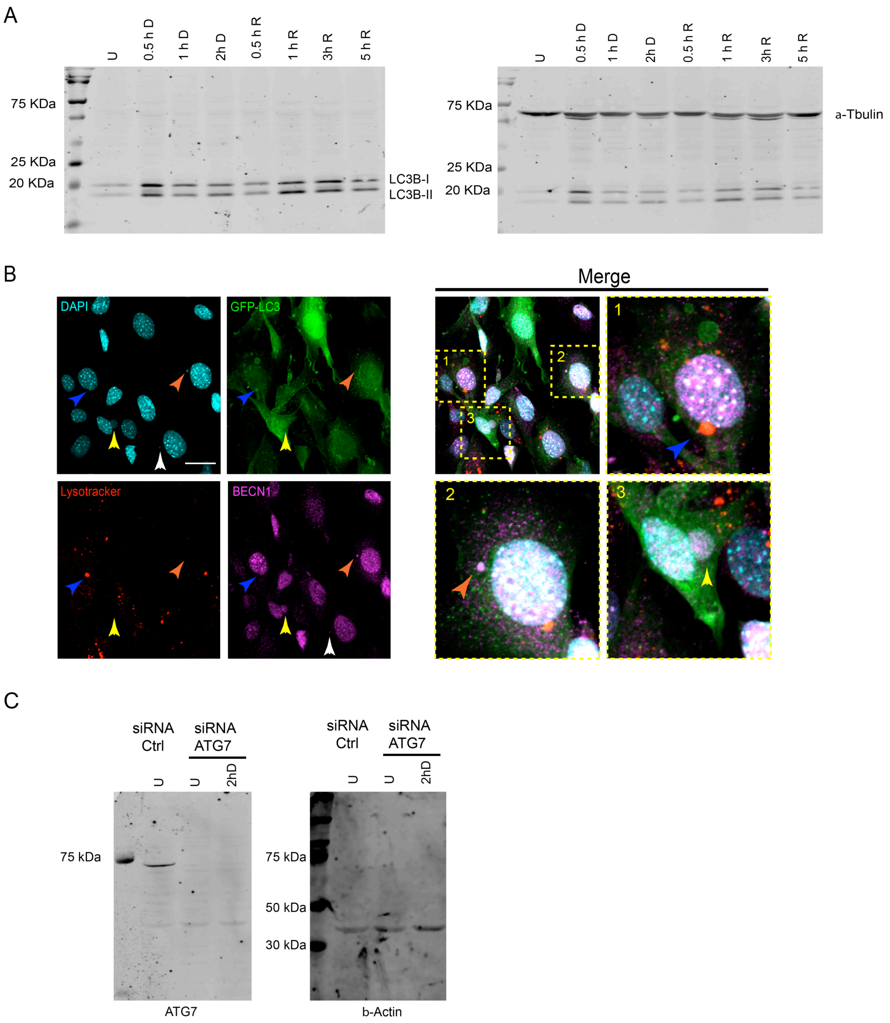


**Figure S2. A.** Whole blots of the WB shown in figure 2A to detect LC3-I, LC3-II and $\alpha$-Tubulin. **B.** Some micronuclei look surrounded by autophagic markers and acidic vesicles. Orange arrows indicate co-localization between DNA and BECN1, yellow arrows show co-localization between DNA, GFP-LC3 and BECN1, and blue arrows indicate a micronucleus surrounded by GFP-LC3 and Lysotracker® (autolysosome). Yellow squares correspond to amplified sections. Cells were treated with Etoposide for 2h. Scale bars correspond to 30 $\mu$m. **C.** Whole membranes for WB to detect ATG7 shown in Figure 2I. MEFs were transfected with siRNA-Atg7 or control siRNA for 48h and then untreated or treated with Etoposide for 2h to detect before total protein extraction. β-actin was detected as loading control.

FIGURE S3

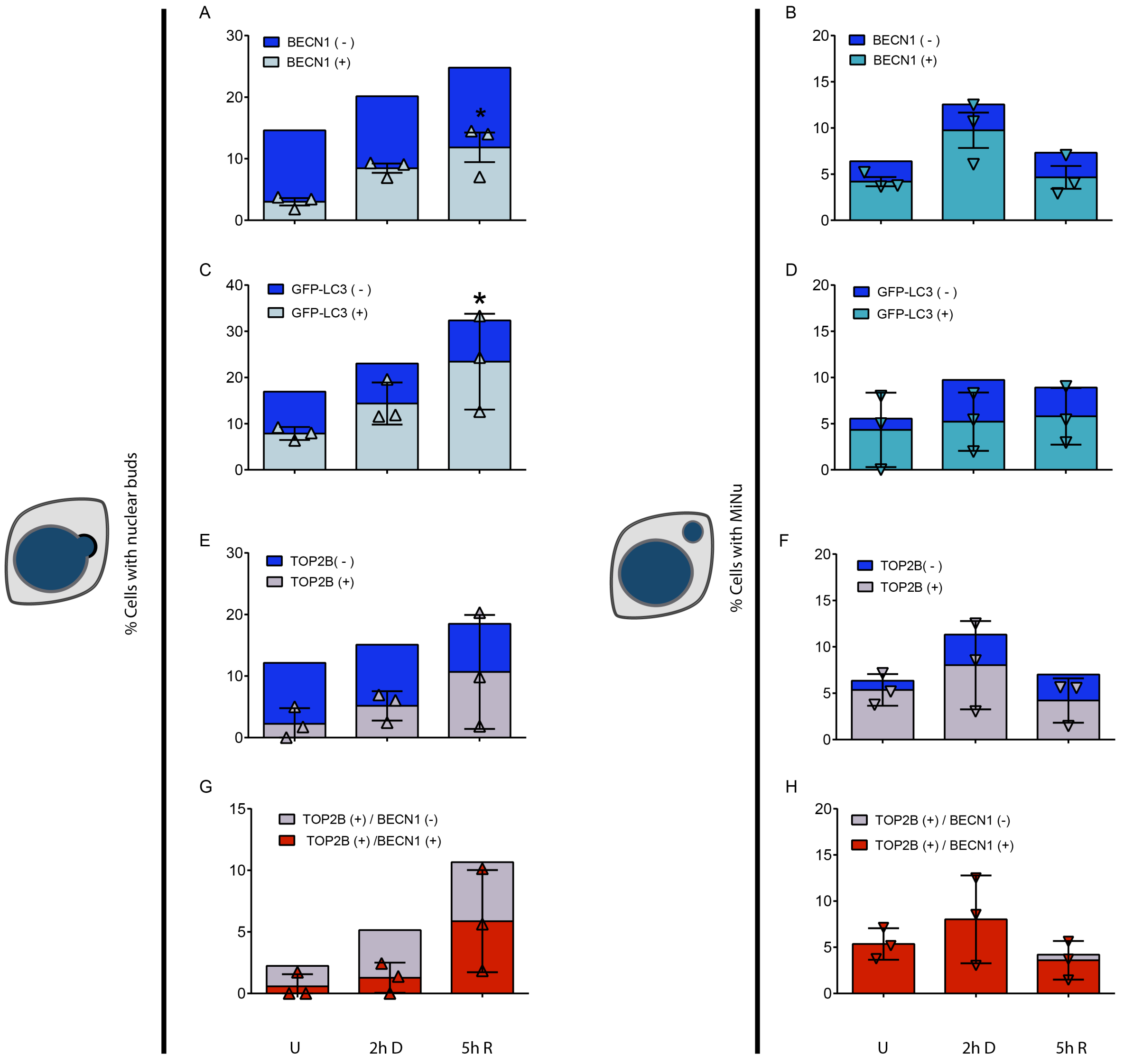

**Figure S3. Nuclear alterations contain autophagic markers and TOP2B. A, B, C and D.** Complimentary graphs showing the differential quantification of nuclear buds and micronuclei containing Beclin-1 (figures **A** and **B**) or LC3 (GFP-LC3 in figures **C** and **D**) or TOP2B (figures **E** and **F**). **G and H**. Complimentary quantifications of nuclear alterations containing simultaneously Beclin-1 and TOP2B. The mean of three independent experiments are graphed. Only for the indicated proteins are presented the result for every experiment as symbols (triangles and inverted triangles) with bars representing SD.  Just for buds containing GFP-LC3, (*) $p<0.05$, Kruskal-Wallis test followed by Dunn's multiple comparison test.
